# Prevalence of Refractive Error and Need for Corrective Lenses in Medically Underserved Residents of Tijuana, Mexico

**DOI:** 10.1101/602805

**Authors:** John L. Ubels, Jonathan M. Ismond, Micah A. Timmermans, Arlene J. Hoogewerf

**Affiliations:** Department of Biology, Calvin College, Grand Rapids, MI USA; The Public Health Program, Calvin College, Grand Rapids, MI USA

## Abstract

**Purpose:** The population of Tijuana, Mexico is growing rapidly, with a current official population estimate of 1.7 million. Nearly 80,000 people migrate to Tijuana each year, resulting in the rise of neighborhoods with substandard housing, lack of services and inadequate access to health care, including eye care. This study describes refractive errors and the need for corrective lenses among participants attending free clinics in these neighborhoods where they received free eye exams and glasses during January 2016. *Methods*: This is a retrospective observational chart review of de-identified data collected from intake forms that were filled out for each participant at the clinics. Subjects were self-selected in response to announcements in the neighborhoods where clinics were conducted. Subjects with presenting uncorrected visual acuity 20/30 OU or worse were examined with an autorefractor to measure spherical refractive error. Either prescription or reading glasses were then distributed to participants who had refractive errors. Epi Info, an open source program provided by the CDC, was used to analyze demographic, visual acuity and refractive error data. *Results*: Presenting visual acuity was evaluated in 1209 people. Of these patients, 70% had a visual acuity of 20/30 or worse. Only 23% of these patients had glasses. Among the patients who were given refractions, 13% had clinically significant myopia (−0.75 D or worse in at least one eye). In participants 20 years old and younger, only 8% had clinically significant myopia. Clinically significant hyperopia (+0.75 D or worse in at least one eye) was detected in 25% of participants. Astigmatism (−1.5 D or worse in at least one eye) was present in 18% of participants. Prescription glasses were given to 542 participants and 396 of these people received their first glasses. Reading glasses were given to 386 people. Among students only 15% presented at the clinics with glasses, while it was determined that 56% of student participants needed glasses. *Conclusion*: The high levels of uncorrected refractive error in this study suggest limited access to affordable eye care in neighborhoods where clinics were conducted. Prevalence of myopia among adolescents and young adults is increasing in many parts of the world. In contrast, a relatively high prevalence of hyperopia was observed in this age group in Tijuana. The data demonstrate an urgent need for eye care and correction of refractive error in the study group.

## Introduction

According to the World Health Organization, uncorrected refractive error is the leading cause of visual impairment globally. Of the 285 million people who are visually impaired, over 80% have preventable visual impairment, 42% due to uncorrected refractive error and 33% due to cataract^1^ (https://www.who.int/blindness/actionplan/en/; accessed March 25, 2019) This uncorrected refractive error is a major public health problem in developing countries, as well as among socioeconomically disadvantaged groups in the United States.^2^ Inadequate eye care and income that is insufficient to pay for glasses, leads to low quality of life, delayed educational progress by students, billions of dollars of lost economic productivity^3,4^ and impaired driving (https://www.nytimes.com/2018/05/05/health/glasses-developing-world-global-health.html,; https://viiproduction.s3.amazonaws.com/uploads/research_article/pdf/51356f5ddd57fa3f6b000001/VisionImpactInstitute-WhitePaper-Nov12.pdf, accessed March 25, 2019) Interventions in which free glasses are given to people who cannot afford them are known to be effective, as shown by Patel et. al^5^ who distributed reading glasses to presbyopic patients in Tanzania. Providing free glasses to underserved rural children in China has been shown to improve performance in school.^6^

Impoverished neighborhoods in Tijuana, Mexico are an example of a region where uncorrected refractive error is common. Tijuana grew threefold from 1980-2010, with an official population over 1.7 million in 2019 (http://worldpopulationreview.com/world-cities/tijuana-population/, accessed March 25, 2019), while the actual population is estimated at 2,060,000. This growth is fueled by the migration of nearly 80,000 people to Tijuana each year, resulting in the rise of neighborhoods with substandard housing and lack of services. These residents generally have low income jobs, resulting in inadequate access to health care, including eye care.

To address this lack of access to eye care, students and eye care professionals from Calvin College, Grand Rapids Michigan, USA have conducted clinics in Tijuana providing free eye examinations and used glasses in these underserved neighborhoods. To document the unmet need for correction of refractive error among the participants in these clinics, this report describes the demographics, prevalence of refractive error and the need for corrective lenses in the group of 1209 people who participated in these clinics in January 2016.

Because of ethical considerations, short-term medical mission trips of the type during which the data in this report were collected must be undertaken with caution and careful planning.^7-11^ Mission trips in which free glasses are provided to people with uncorrected refractive error are considered to be effective and to do no harm (http://www.uniteforsight.org/; accessed March 25, 2019). The mission trip during which the data in this paper were collected largely adhered to the recommendations of Suchdev et al.^12^ in that the clinics were conducted in response to a need identified by community leaders and were conducted in collaboration with members of the community. Students were educated about the culture and needs of the community, as well as principles of refractive error, eye examinations and prescribing glasses. A team approach was employed in which the students were closely supervised by eye care professionals and worked closely with community members who helped to coordinate the clinics. Finally, this report serves to evaluate the effectiveness of the clinics.

## Methods

The data presented in this study were collected during clinics conducted in churches located in underserved neighborhoods in Tijuana, Mexico in January 2016. The clinic locations were chosen in consultation with local pastors of Reformed Presbyterian Church of Mexico congregations who were aware of economic conditions in the neighborhoods that lead to lack of health care and particularly access to corrective lenses. Residents of the neighborhoods were made aware of the clinics through audio announcements, posters, distribution of flyers and canvassing by church members and clinic volunteers. The clinics were conducted by an ophthalmologist, optometrist and pre-optometry, pre-medical, public health and nursing students from Calvin College in Grand Rapids, Michigan, USA.

Clinic participants filled out an intake form that collected data on gender, age, occupation and literacy. Also noted was the participant’s chief visual complaint and whether she or he had glasses. Presenting visual acuity was then measured using a Snellen chart. Participants with visual acuity 20/30 or worse were given a non-cycloplegic examination with an autorefractor to determine refractive error and appropriate prescription. They were then sent to dispensing where they received used glasses that best matched their required prescription. Adults and children who tested 20/30 or better but complained of difficulty in reading were tested with a near card and provided with reading glasses. Data on presenting, uncorrected visual acuity, presenting refractive error and reader strength were recorded on the intake forms which were collected when the participant left the clinic.

The glasses provided were used prescription glasses obtained from the Lions Club of Grand Rapids, MI and Robert Huizinga, O.D. of Jenison, MI. Prescriptions of the glasses were determined and the placed in a database arranged in a way that permitted volunteers to conveniently find appropriate glasses for clinic participants. There were about 1200 prescription glasses available in the database. Reading glasses were donated used or new glasses.

This study is a retrospective, observational chart review of de-identified data collected from the intake forms that were filled out for each clinic participant. The study was approved by the Calvin College Institutional Review Board. *Epi Info™*, an open source epidemiology program provided by the US Centers for Disease Control (https://www.cdc.gov/epiinfo/index.html, accessed March 25, 2019), was used to analyze data on demographics, presenting visual acuity, refractive error and need for corrective lenses. Limitations of the study are: 1) clinic participants were self-selected based on their awareness of the clinic announcements and their own perceived need for eye care. Therefore, this not a population-based study; 2) For security reasons, the clinics were held only during the day. This may have made it difficult for employed persons to attend. Students could attend either before or after the school day. 3) Literacy was self-reported. A literacy test was not administered.

## Results and Discussion

### Demographic information

A total of 1209 people participated in the clinics. Females outnumbered males by a ratio of 1.5:1. The preponderance of women was especially evident in the age range of 20-49 years old in which women outnumbered men by 2.6:1 (Table 1). This is in agreement with 415 people identifying their occupation as homemaker (Table 2) and the timing of the clinics which were only held during the day when many men were at work. Relatively few people over the age of 70, (maximum age 86) attended the clinics. This may reflect lower numbers of elderly people in the neighborhoods, for which we have no data, or difficulty for older people in getting to the clinics, since many participants walked.

**Table 1.** Ages of clinic participants*

|  | Males |  | Females |  | Total |  |
| --- | --- | --- | --- | --- | --- | --- |
| Age | n | % | n | % | n | % |
| 1 - 9 | 79 | 59 | 55 | 41 | 134 | 11 |
| 10 - 19 | 93 | 47 | 104 | 53 | 197 | 16 |
| 20 - 29 | 23 | 23 | 76 | 77 | 99 | 8 |
| 30 - 39 | 33 | 24 | 104 | 76 | 137 | 11 |
| 40 - 49 | 73 | 31 | 159 | 69 | 232 | 19 |
| 50 - 59 | 87 | 41 | 126 | 59 | 213 | 18 |
| 60 - 69 | 49 | 45 | 60 | 55 | 109 | 9 |
| ≥ 70 | 36 | 47 | 40 | 53 | 76 | 6 |
| Total | 473 | 40 | 724 | 60 | 1197 |  |
| Unknown age or gender: 12 |  |  |  |  |  |  |
|  | Mean Age | Std Dev | Median Age |  |  |  |
| Male | 36.7 | 23.0 | 41.0 |  |  |  |
| Female | 38.9 | 19.5 | 41.0 |  |  |  |

\*This table includes all clinic participants, a total of 1209. Data in subsequent tables may not add up to this total due to missing data on intake forms.

**Table 2.** Occupations of clinic participants with the number of participants in each category.

|  |  |
| --- | --- |
| Homemaker | 415 |
| Child - pre-school (1-4 years old) | 7 |
| Student/school age | 321 |
| Factory worker | 165 |
| Vendor/sales | 47 |
| Construction/building trades | 44 |
| Mechanic | 19 |
| Service industries (various) | 17 |
| Retired/disabled | 17 |
| Driver (taxi, truck) | 16 |
| Pastor/church worker | 15 |
| Security | 12 |
| Custodial/maintenance/housekeeping | 11 |
| Restaurant/food industry | 10 |
| Administrative/office work | 8 |
| Other | 5 |
| Unemployed | 27 |
| Unknown | 83 |

**Table 3.**
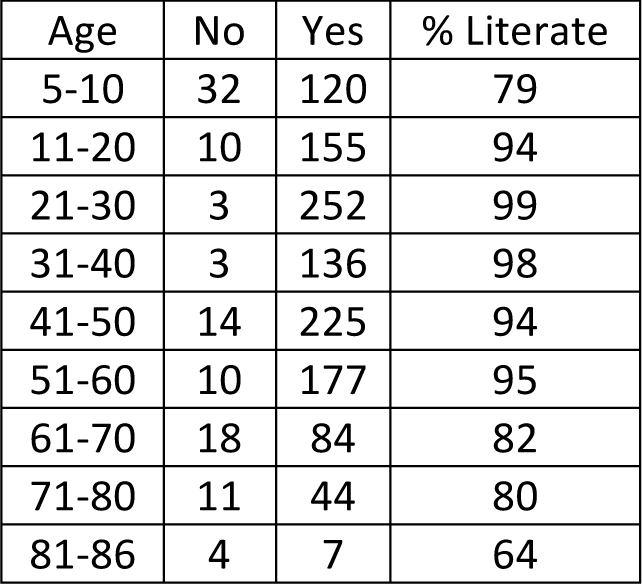
Self-reported literacy rates among clinic participants.

| Age | No | Yes | % Literate |
| --- | --- | --- | --- |
| 5-10 | 32 | 120 | 79 |
| 11-20 | 10 | 155 | 94 |
| 21-30 | 3 | 252 | 99 |
| 31-40 | 3 | 136 | 98 |
| 41-50 | 14 | 225 | 94 |
| 51-60 | 10 | 177 | 95 |
| 61-70 | 18 | 84 | 82 |
| 71-80 | 11 | 44 | 80 |
| 81-86 | 4 | 7 | 64 |

The most common occupation among clinic participants was homemaker, reflecting the location of the clinics in neighborhoods and, as stated above, the timing of the clinics between 9 AM and 4 PM (Table 2). A large number of school age children were brought to the clinics by their mothers, which included 284 who identified as elementary to high school age students. Only 12 identified themselves as post-high school age students (over 18 years old). The number of participants in the school age category (321) is greater than the number of students because some children do not go to school. According to the pastors of the churches where the clinics were held, although a free public education is guaranteed through the eighth grade, some children do not go to school because their parents cannot afford the added costs of books and uniforms. Admission to high school is based on an entrance test.

Reflecting the demographics of the neighborhoods, most occupations are in the factories and service industries. The most frequent occupation was factory worker, in keeping with the presence in the neighborhoods of large maquiladoras (assembly plants) which are the primary industry in Tijuana. If it had been possible to hold clinics at night, it is expected that it would have been possible to provide eye exams to a larger number of employed people. The large number of participants identifying themselves as vendors reflects the presence of street markets near several of the clinics.

While 396 clinic participants identified themselves as having a job, only 27 self-identified as unemployed. At the time of the clinics the official unemployment rate in Baja California, where Tijuana is located, was 3.6% (http://www.thebajapost.com/2015/12/08/record-unemployment-rate-figures-in-baja-california/; http://www.bajacalifornia.gob.mx/sedeco/, accessed March 25, 2019) with a labor force participation rate of about 60% (https://tradingeconomics.com/mexico/labor-force-participation-rate, accessed March 25, 2019). It should be noted that some of the clinic participants who did not report an occupation (“Unknown” in Table 2) may have been unemployed, retired or not participating in the labor force.

The self-reported literacy rate of clinic participants between the ages of 11 and 86 years old was 94%. This agrees with the reported literacy rate of 95% in Mexico (https://www.cia.gov/library/publications/the-world-factbook/fields/370.html; accessed March 25, 2019). These data provide evidence for the importance of providing prescription lenses and reading glasses to the members of the study group who have refractive error but apparently cannot afford glasses.

### Presenting Visual Acuity

The presenting, uncorrected visual acuity of the clinic participants is shown in Tables 4 and 5. It must be emphasized that these individuals do not represent a random sample of the people living in the neighborhoods where the clinics were held. Rather these are people who were aware of the clinics and attended because of a perceived need for glasses or a desire to have an eye exam for themselves or their children. Therefore, comparisons of the data in this paper to published, population-based studies from other regions must be done with the caveat that there were people in the neighborhoods who did not need glasses, were able to afford glasses or were not able to attend the clinics.

**Table 4.** Uncorrected presenting visual acuity, by age group. Participants with acuity 20/30 or worse received refractions.

| Age | Number of individuals | Better than 20/30 OU | 20/30 or worse OU | % 20/30 or worse OU |
| --- | --- | --- | --- | --- |
| 1-20 | 342 | 164 | 178 | 52 |
| 21-40 | 252 | 131 | 121 | 48 |
| >40 | 615 | 65 | 550 | 89 |
| Total | 1209 | 360 | 849 | 70 |

**Table 5.** Distribution of uncorrected presenting visual acuity.

| Visual Acuity | OD |  | OS |  | OU |  |
| --- | --- | --- | --- | --- | --- | --- |
|  | Number of Individuals* | % | Number of Individuals | % | Number of Individuals | % |
| 20/20 or better | 210 | 18.3 | 222 | 19.4 | 216 | 19.1 |
| 20/25 to 20/30 | 377 | 32.9 | 401 | 35.1 | 305 | 27.0 |
| 20/40 to 20/50 | 221 | 19.2 | 211 | 18.4 | 299 | 26.5 |
| 20/60 to 20/70 | 120 | 10.5 | 122 | 10.7 | 95 | 8.4 |
| 20/80 to 20/100 | 92 | 8.0 | 67 | 5.9 | 199 | 17.6 |
| 20/200 or worse | 127 | 11.1 | 121 | 10.6 | 14 | 1.2 |
| Counting fingers | 27 |  | 17 |  | 5 |  |
| Cannot count fingers | 8 |  | 7 |  | 3 |  |

\*Totals do not add up to total clinic participants because of missing data. Visual acuity was not recorded for 55 individuals. Of these, 12 did not receive glasses, 31 received only reading glasses suggesting that their visual acuity was better than 20/30, and the remaining 12 received prescription glasses.

In the age group 40 years or younger about 50% participants had a presenting visual acuity of 20/30 or worse. Of those over 40, 89% needed prescription lenses. The cut off of 20/30 for further examination was based on the limited availability of lower power lenses in the supply of used glasses. In total, 70% of clinic participants were sent for refraction due to a presenting visual acuity of 20/30 or worse. To place these numbers in context, 32% of Americans over 40 wear corrective lenses for myopia or hyperopia and 46% of the entire population uses corrective lenses (https://www.aao.org/newsroom/eye-health-statistics, accessed March 25, 2019). In Europe about 50% of adults 25 – 90 years old have refractive error.^13^

As shown in Table 5, some clinic participants had uncorrectable visual impairment, commonly due to cataracts, pterygium impinging on the visual axis, corneal scars and in a few cases, retinal disease. These people were referred to the pastors, when possible, for assistance in seeking care from an ophthalmologist.

### Prevalence of Myopia

Myopia, defined in this study as a measured sphere of refractive error of −0.75 D or worse in either eye, was uncommon among the clinic participants. While there is currently an epidemic of myopia among children and young adults in many countries,^14^ only 8% of clinic participants 20 years old and younger were myopic (Table 6; Fig. 1). This contrasts with a population based study in Monterey, Mexico that reported myopia in 27% of 12 and 13 year olds.^15^ Gomez-Salazar et al.^16^ conducted a study of refractive error in nearly 700,000 patients, age 6-90, visiting primary care optometry clinics for low income patients in 14 Mexican states and reported a 25% prevalence of myopia nationwide and 27% in Baja California, the location of Tijuana. This is similar to the 27% prevalence of myopia in clinic participants aged 21-40 in the present study (Table 6). In another Hispanic country, Paraguay the prevalence of myopia was 38% in 3-22 year olds.^17^ Prevalence of myopia among adults in the US and Europe is reported to be 25-33% and 37%, respectively.^13,18,19^ In contrast, the prevalence of myopia among children and young adults exceeds 70% in several Asian cities, with 97% of 19 year-old men in Seoul, Korea being myopic.^20^

**Table 6.**
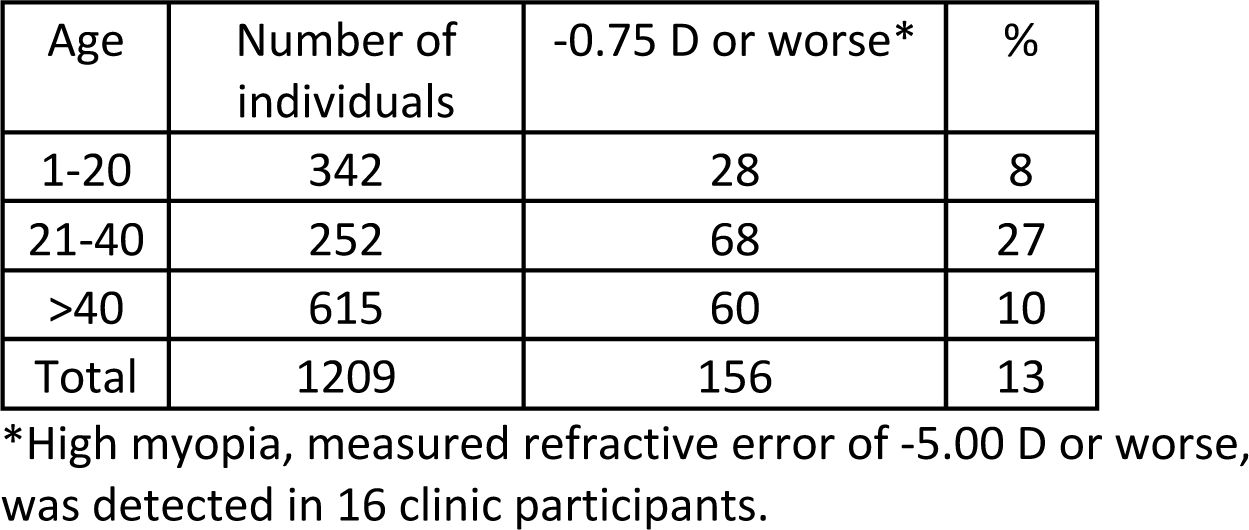
Prevalence of myopia in either eye, by age group.

**Fig. 1.**
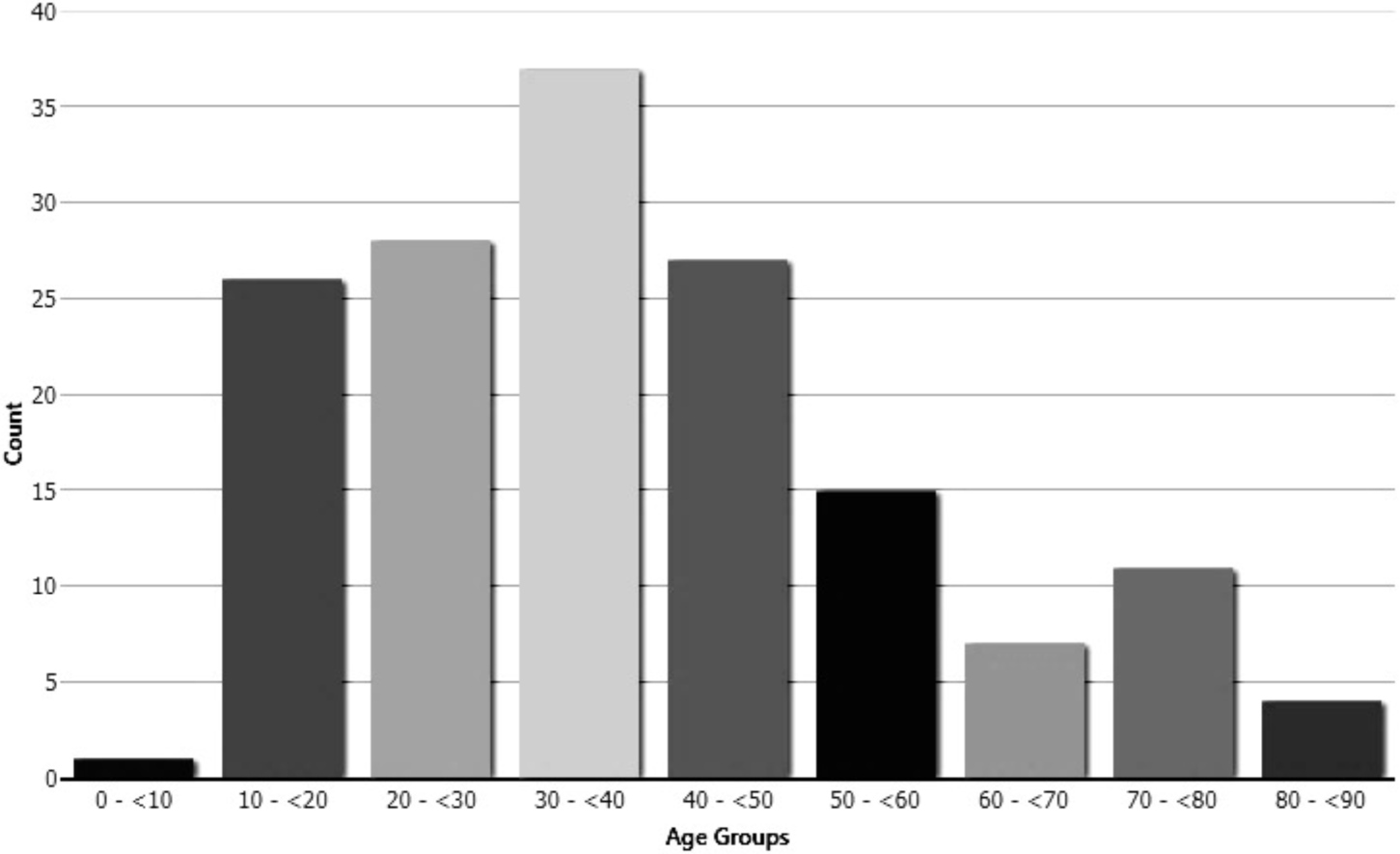
Numbers of clinic participants with myopia −0.75 D or worse, by decade. The prevalence of myopia is relatively low in the age group 10-20 years old.

The relatively low prevalence of myopia among children and young adults from the Tijauna neighborhoods who attended our clinics is of interest. In the Gomez-Salazar et al.^16^ report, 20% of children 6-9 years old and 41% of those 10-19 years old were myopic. The Gomez-Salazar study may be compared to the present study, because the patients were self-selected, based on a decision to be examined at an optometry clinic. For many years, it was thought that near work, such as reading, was a risk factor for myopia. It is now recognized that lack of light exposure is a risk factor for myopia and that increased time spent outdoors is correlated with lower prevalence of myopia.^20-25^ It may be that higher socioeconomic status results in more time indoors because of the opportunity to spend time in reading and computer use. A study in Nepal reported myopia in 27% of upper-middle class 15 year old students attending elite schools Kathmandu compared to 3% in 15 year olds in a rural district with low school attendance rates and greater time spent outdoors.^26^ Gomez-Salazar et al.^16^ reported myopia prevalence of only 19% in the Mexican states of Nayarit and Sinaloa which are rural and agricultural. The neighborhoods where our clinics were held are urban and most children attend school. A study of time outdoors in these neighborhoods would be of interest, since the children only attend school in the morning or afternoon, and anecdotally, we have observed that they live in very small houses and play outdoors frequently.

### Prevalence of Hyperopia

In this study hyperopia was defined as measured sphere of refractive error of +0.75 D or worse and was observed in 25% of clinic participants. Moderate to high hyperopia, measured refractive error of +3.00 D or worse, was detected in 78 people or 6.5% of the clinic participants. This may be compared to a 9% prevalence of refractive error of +3.00 D or worse among Hispanics in the US in a population based study.^19^ There was, however, great variability in the prevalence of hyperopia among age groups (Table 6; Fig. 2). While, as discussed above, myopia was less common than expected in children and young adults, 24% of clinic participants 20 years old and younger were hyperopic.

**Table 6.** Prevalence of hyperopia in either eye, by age group.

| Age | Number of individuals | +0.75 D or worse | % |
| --- | --- | --- | --- |
| 0-20 | 342 | 83 | 24.3% |
| 21-40 | 252 | 11 | 4.4% |
| >40 | 615 | 206 | 33.5% |

**Fig. 2.**
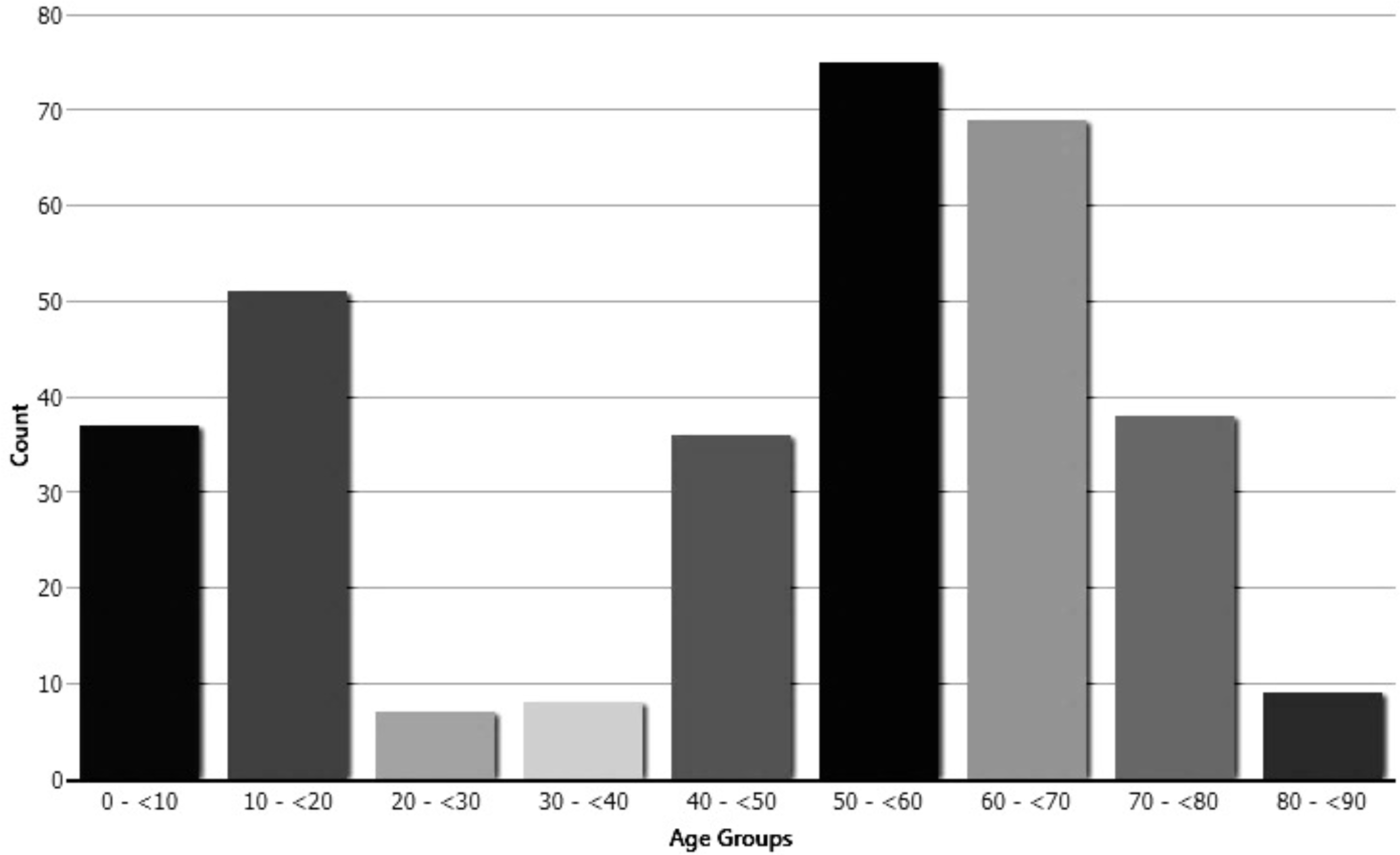
Numbers of clinic participants with hyperopia +0.75 D or worse, by decade.

With the caveat that this is not a population based study, we place these data in the context of previous reports on hyperopia. It is well known that hyperopia occurs commonly in young children and decreases in prevalence with age. Castagno et al.^27^ reported an 8% prevalence of hyperopia in Brazilian 6 year olds, decreasing to 1% at age 15. Gomez-Salazar et al.^16^ reported a similar prevalence and trend to decreased hyperopia in 6-19 year olds in Mexico. Prevalence of hyperopia was higher in this age group in the present study.

This prevalence of hyperopia decreased dramatically to 4.4% in the 21-40 year old age group. This is agreement with the prevalence of hyperopia in this age group in the Gomez-Salazar et al.^16^ study. In keeping with the hyperopic shift known to occur in older people, ^28-30^ hyperopia increased to 34% in participants over 40, a similar prevalence to that reported in Mexican patients by Gomez-Salazar et al.^16^ Prevalence of hyperopia in people over 40 has been reported to be 25% in population based studies in the US and Europe.^13,19^

### Astigmatism

Astigmatism was common among clinic participants, with 18% of all participants having astigmatism of −1.50 D or worse (Table 7.). The prevalence by age group agrees with previously reported data^16,17,28,30^ and overall prevalence is similar to that reported by Gomez-Salazar et al.^16^ in Baja California. This may be compared to a prevalence of astigmatism of 23% in all those aged 20-39 in the US and 20% of Mexican Americans in the same age group. An interesting difference is that only 18% of those over 40 years old in the present study had astigmatism while the prevalence in this age group is 31% in the US.^18^

**Table 7.**
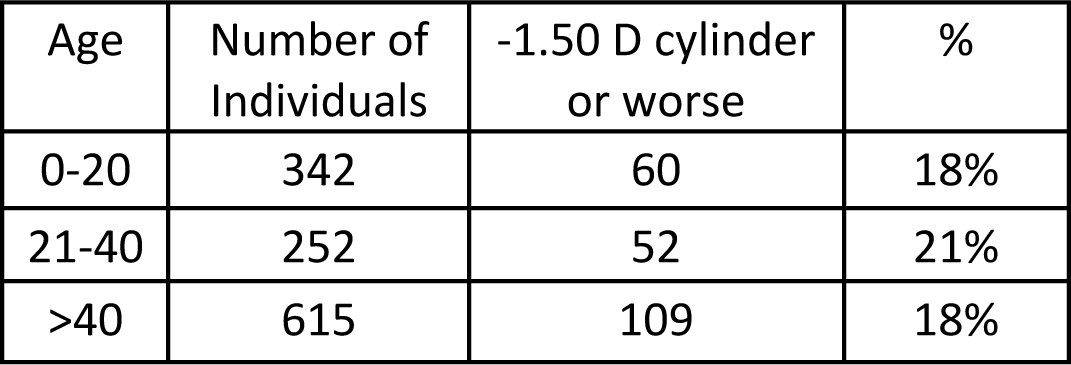
Prevalence of astigmatism in either eye, by age group.

Extreme astigmatism, −4.00 D of cylinder or worse in either eye, was detected in 32 clinic participants (Table 8). These patients were difficult to fit with glasses, since only 1% of the glasses in the database had a cylinder of at least −4.00 D in either lens. We were unable to provide glasses for 4 of them (12.5%). This suggests that extreme astigmatism is relatively rare in the midwestern US population that donated the used glasses. In contrast, of the 556 clinic participants who needed prescription lenses for any reason and had correctable refractive error, only 2.5% did not receive glasses.

**Table 8.** Prevalence of moderate, severe and extreme astigmatism in either eye.

| D | Number of Individuals | % of clinic participants |
| --- | --- | --- |
| Moderate: -1.00 to -1.75 | 216 | 17.9% |
| Severe: -2.00 to -3.75 | 105 | 8.7% |
| Extreme: -4.00 or worse | 32 | 2.6% |

### Clinic Participants Receiving Glasses

The primary goal of the mission trip during which the data in this study were collected was to provide glasses to those in need, and it is apparent that there was a high level of uncorrected refractive error among the clinic participants. Only 23% of participants had prescription glasses prior to coming to the clinic and 45% of participants received glasses. Of those who had glasses, 85% needed new glasses, and 73% of all those who received prescription glasses were given their first pair (Table 9). The groups most frequently requiring prescription glasses were in the age groups 10-19 years old and 40-69 years old. (Fig. 3)

**Table 9.** Number of clinic participants who presented wearing glasses and number of prescription glasses provided.

|  | No | Yes | % Yes |
| --- | --- | --- | --- |
| Participants with glasses prior to clinic | 936 | 272 | 23% |
| Rx glasses provided to clinic participants | 667 | 542 | 45% |
| Patients with prior glasses receiving replacements | 40 | 232 | 85% |
| Participants receiving first glasses | 145 | 396 | 73% |

**Figure 3.**
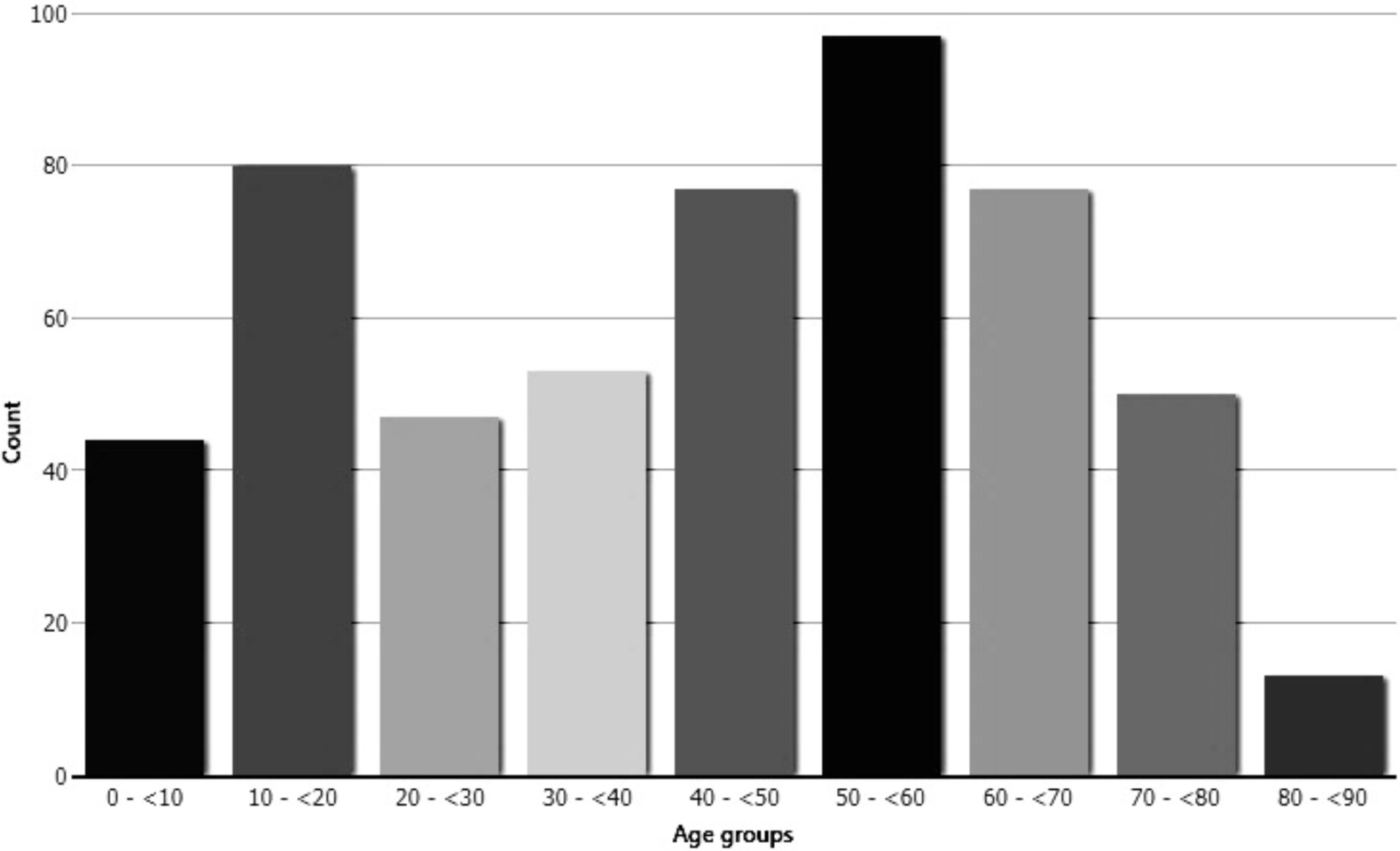
Number of clinic participants receiving prescription glasses by decade.

Because of the importance of good vision to educational success, the students under age 20 were separated for further analysis. Only 15% of the students presenting at the clinics had glasses, and of the 85% who did not have glasses, 41% received glasses. Of those students who had glasses, 61% needed new glasses (Table 10). As noted above, prevalence of myopia was unexpectedly low in this group, and the majority of students with refractive error needed a correction for hyperopia or reading. The numbers of individuals in Table 10 do not match those in Tables 5 and 6 because, due to the importance of good vision, some students with a presenting visual acuity of 20/25 were sent for autorefraction and were given −0.50 or +0.5 glasses, Also, some students with good presenting visual acuity were given plus prescription glasses for reading. Also, some students did not meet the criteria for being myopic or hyperopic, but needed correction for astigmatism. The importance of intervening by providing free glasses to the student is supported by the study of Ma et al.^6^ in China and a recent study in the United States which reports that standardized test scores increased when, in addition to vision screening, free eye exams and free glasses were provided to low-income students.^31^

**Table 10.** Need for glasses by students under 20 years old presenting at the clinics*

|  |  | % |
| --- | --- | --- |
| Total number of students | 296 |  |
| Students presenting without glasses | 252 | 85 |
| Students without glasses who received glasses | 122 | 41 |
| Students presenting with glasses* | 44 | 15 |
| Students with glasses receiving new glasses* | 27 | 61 |
| Students needing prescription glasses | 139 | 47 |
| Students needing reading glasses | 27 | 9 |

\*includes prescription and reading glasses

There was an urgent need for reading glasses among the clinic participants, and 396 pairs were distributed. As expected, because of the known prevalence of presbyopia in this age group, 47% of clinic participants age 30-59 were given reading glasses (Fig. 4). Also, 83% of the participants in this age group who received prescription glasses needed bifocals or progressive lenses for reading.

**Figure 4.**
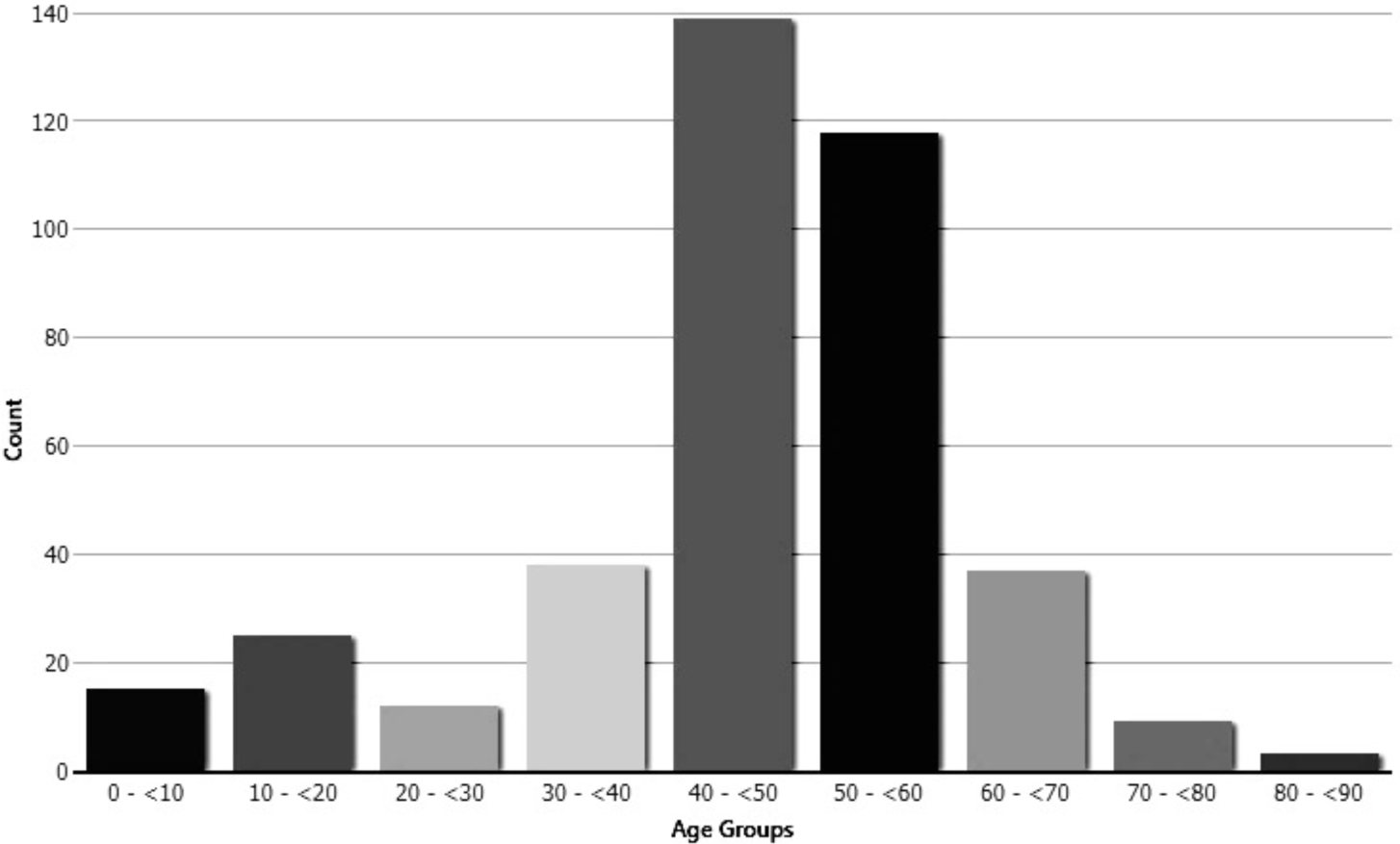
Number of clinic participants receiving reading glasses, by decade. 53.4% of glasses provided were in the range of +1.50 to +2.00 D.

The lack of glasses among clinic participants with uncorrected refractive error, despite high rates of literacy and employment, suggests that the residents of the neighborhoods where the clinics were held are using limited resources for necessities such as food, clothing and shelter, leaving inadequate funds for eye care. This problem can be understood in the context of a review of wages in Mexico as compared to the cost of glasses. All values are expressed in US dollars. Although the minimum wage in Mexico has recently increased to $4.70/hour, at the time of when the clinics in this study were conducted wages were about $2.00/ hour. Factory workers earned about $17/day ($340/month), low skilled workers, $11/day ($220/month) and a living income for a family was $378/month. (https://tradingeconomics.com/mexico/wages-inmanufacturing; accessed March 25, 2019). A perusal of websites of opticians shows that single vision prescription glasses can be purchased in Mexico for about $50, while progressive lenses, which were needed by many people who visited our clinics, cost about $140. Based on employment data in this report (Table 2) showing that many people have low income jobs, this is a significant percentage of monthly income for many of our clinic participants. We observed in stores in Tijuana that reading glasses cost $7-9, nearly a day’s wage for many people. For comparison, in the US, where a factory worker in a similar type of job to factory work in Tijuana makes $2000-3000/month and the minimum wage is $7.25/hr or more, single vision glasses can be obtained at discount optical shops for as little as $70 with a free eye exam. Reading glasses in the US typically cost $4-10 and can be obtained a “dollar” stores, which do not exist in Mexico, for as little as $1. Since prescription glasses in Mexico cost about as much as they do in the US and reading glasses cost more, while wages are much lower, the reason for the high level of uncorrected refractive error is easily understandable.

## Conclusions

Evaluation of data collected during the eye care mission trip described in this report documents a high prevalence of need for corrective lenses in the economically challenged neighborhoods of Tijuana, Mexico where the clinics were conducted. The data also may be compared to prevalence of refractive error in other regions of the world and therefore are of potential value to epidemiologists. In the age group 20 years old and younger in this study prevalence of myopia is low compared to the reported epidemic of myopia in other regions of the world. In contrast, in the age group 20 years old and younger the prevalence of hyperopia is higher than expected, while the prevalence of hyperopia is consistent with other reports for people over age 40. The high need for reading glasses and prescription glasses with bifocals or progressive lenses shows that access to corrective lenses is very important in this highly literate population, many of whom work in assembly plants where good vision is important to the type of work that they do.

The high prevalence of uncorrected visual impairment among the clinic participants suggests limited access to affordable eye care in the neighborhoods where the clinics were conducted. It is important that the work of short-term mission trips be sustainable and that the root cause of problems be addressed.^8,12^ The data in this report are from one of four eye care trips that have been conducted by our group to the neighborhoods described over a period of 10 years. Economic conditions and practices in Tijuana are such that it is difficult to directly address the root cause of inability to afford glasses. As such, interventions of the type described in this report continue to be important.

## Declarations

This study was approved by the Calvin College Institutional Review Board. Consent to participate was not required for this study.

## Acknowledgements

We thank the pastors and members of the Iglesia Presbiteriana Reformada de Mexico and the missionaries and staff of the Christian Reformed Church in North America for helping to organize the clinics. Larry Gerbens, M.D., Ralph De Haan, O.D. and John L. Ubels, Ph.D. supervised the clinics. The Lions Club of Grand Rapids, MI and Robert Huizenga, O.D. provided used glasses, and students in the Public Health 248 Epidemiology course at Calvin College assisted with data analysis under the supervision of Arlene Hoogewerf, Ph.D. The authors also thank Brittany Hoolsema, O.D., for providing a review on the ethics and effectiveness of short-term missions.

Supported by the Calvin College Fund for Eye Research, West Michigan Optometric Scholarship (JMI), the Wagner Memorial Scholarship for pre-optometry students (MAT) and a gift to Calvin College from Robert and Anita Huizenga.

